# Decoupled Maternal and Zygotic Genetic Effects Shape the Evolution of Development

**DOI:** 10.1101/200774

**Authors:** Christina Zakas, Jennifer M. Deutscher, Alex D. Kay, Matthew V. Rockman

## Abstract

Many animals develop indirectly via a larval stage that is morphologically and ecologically distinct from its adult form. Hundreds of lineages across animal phylogeny have secondarily lost larval forms, instead producing offspring that directly develop into adult form without a distinct larval ecological niche^1–7^. Indirect development in the sea is typically planktotrophic: females produce large numbers of small offspring that require exogenous planktonic food to develop before metamorphosing into benthic juveniles. Direct development is typically lecithotrophic: females produce a smaller number of larger eggs, each developing into a juvenile without the need for larval feeding, provisioned by yolk. Evolutionary theory suggests that these alternative developmental strategies represent stable alternative fitness peaks, while intermediate states are disfavored^4,8–11^. Transitions from planktotrophy to lecithotrophy thus require crossing a fitness valley and represent radical and coordinated transformations of life-history, fecundity, ecology, dispersal, and development^7,12–16^. Here we dissect this transition in *Streblospio benedicti*, the sole genetically tractable species that harbors both states as heritable variation^17–19^. We identify large-effect loci that act maternally to influence larval size and independent, unlinked large-effect loci that act zygotically to affect discrete aspects of larval morphology. Because lecithotrophs and planktotrophs differ in both size and morphology, the genetic basis of larval form exhibits strong maternal-by-zygotic epistasis for fitness^20^. The fitness of zygotic alleles depends on their maternal background, creating a positive frequency-dependence that may homogenize local populations. Developmental and population genetics interact to shape larval evolution.

---

*Streblospio benedicti* females fall into two classes exhibiting classic life-history tradeoffs: Planktotrophic mothers produce small (~100um) eggs that develop into obligately feeding larvae that spend weeks in the plankton. These larvae are pelagic and grow larva-specific swimming chaetae that are thought to deter predation^21^ (Fig 1a). Lecithotrophic mothers produce large (~200um) eggs that develop into larvae that do not require exogenous food and quickly leave the water column as benthic juveniles. These larvae lack swimming chaetae and a second type of larva-specific morphological structure, anal cirri containing bacillary cells, which are distinctive rhabdomeric cells of unknown function^22^ (Fig 1a). There are also differences during embryogenesis^23^ and organogenesis^22,24^, where accelerated development of juvenile features and truncated development of larval features occurs in lecithotrophs. Despite these developmental and larval differences, adults are indistinguishable at the level of gross morphology. Unlike examples of polyphenism, environmental input does not substantially alter egg size or subsequent offspring type^25,26^. *S. benedicti* is common throughout North American estuaries, but local populations are usually uniformly one larval type^17,27,28^. The larval mode in the population does not correlate with any known environmental gradient. Despite the persistence of these two dramatically different strategies in nature, they canbe crossed in the lab producing an array of morphological and developmental intermediates (Fig 1a).

**Figure 1.**
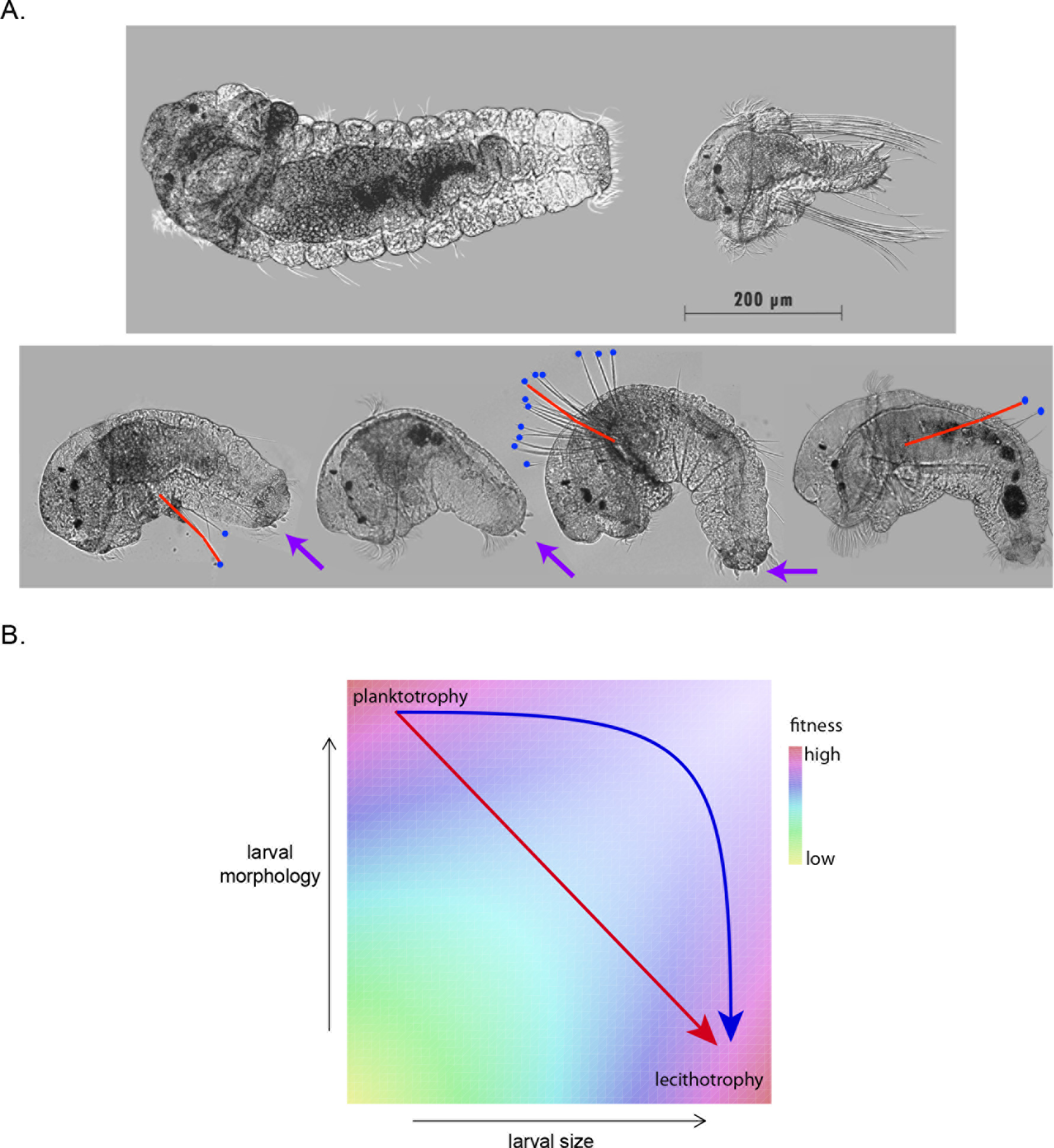
*S. benedicti* larval morphology. **A.** Wild populations occupy two extremes of larval size and form: planktotrophic larvae (top right) carry larva-specific chaetae and anal cirri, while lecithotrophic larvae (top left) lack these traits and are capable of adopting a benthic habit without feeding. G_2_ animals (second row) are intermediate in size and variable in their larval morphologies. We measured larval area, chaetae length (red), presence of anal cirri (purple arrows) and number of chaetae per side, when present (blue dots). **B.** Evolutionary theory and the absence of intermediates in the wild suggest a hypothetical fitness landscape, in which planktotrophs and lecithotrophs occupy fitness peaks separated by a valley in size-shape-fitness space. Transitions from planktotrophy to lecithotrophy may involve single-step pleiotropic transformations (red arrow) or independent evolution of size and form (blue arrow). The latter scenario entails producing an intermediate larval type with a reduction in fitness.

Genetic models for evolutionary transitions from planktotrophy to lecithotrophy require traversal of a fitness valley in multivariate phenotype space^29^ (Fig 1b). The simplest scenario is that co-adapted trait combinations are affected by shared alleles, either by pleiotropy (for example, ref ^30^) or tight linkage (e.g., a supergene with suppressed recombination^31,32^). The necessity to coordinate maternal traits (larval size) with zygotic traits (larval morphology) adds additional constraints^20,33,34^. Fig 1b shows that alternative paths from planktotrophy to lecithotrophy make different predictions about the correlations among genetic effects. In particular, the presence of facultative planktotrophy (i.e., larvae can feed but do not require food) in lecithotrophic *S. benedicti* larvae^35,36^ provides one suggestion that evolution could traverse the larval-size axis, from small to large, prior to traversing the larval morphology axis, from ‘complex’ feeding larva to ‘simple’ non-feeding larvae with accelerated development of juvenile features^3,6^. Nevertheless, in natural populations we observe only the two extremes: large, ‘simple’ lecithotrophs, and small, ‘complex’ planktotrophs.

We dissected developmental mode by crossing a lecithotrophic male and a planktotrophic female, each derived from a well-characterized homogeneous population^19,37,38^. From the F_1_ progeny of this cross between a Long Beach (CA) lecithotroph and Bayonne (NJ) planktotroph, we generated a G_2_ panel by crossing one F_1_ male with four F_1_ females. We measured larval features - size, number and length of swimming chaetae, and presence or absence of anal cirri - on each G_2_ animal, then raised it to adulthood and crossed it to another G_2_ animal. From the progeny of these crosses, we measured average size of each G_2_ female’s G_3_ offspring. This extra generation is required because larval size is a maternal-effect trait^19^. Indeed, we find that the larval size of a G_2_ female is uncorrelated with the average size of her G_3_ progeny (Fig 2a). The G_2_ larvae all develop with equivalent maternal genetic effects, as their F_1_ mothers are heterozygotes for planktotrophic and lecithotrophic alleles. By contrast, maternal effect genes are segregating among G_2_ females, and so G_3_ brood sizes reflect inherited maternal-effect variation.

**Figure 2.**
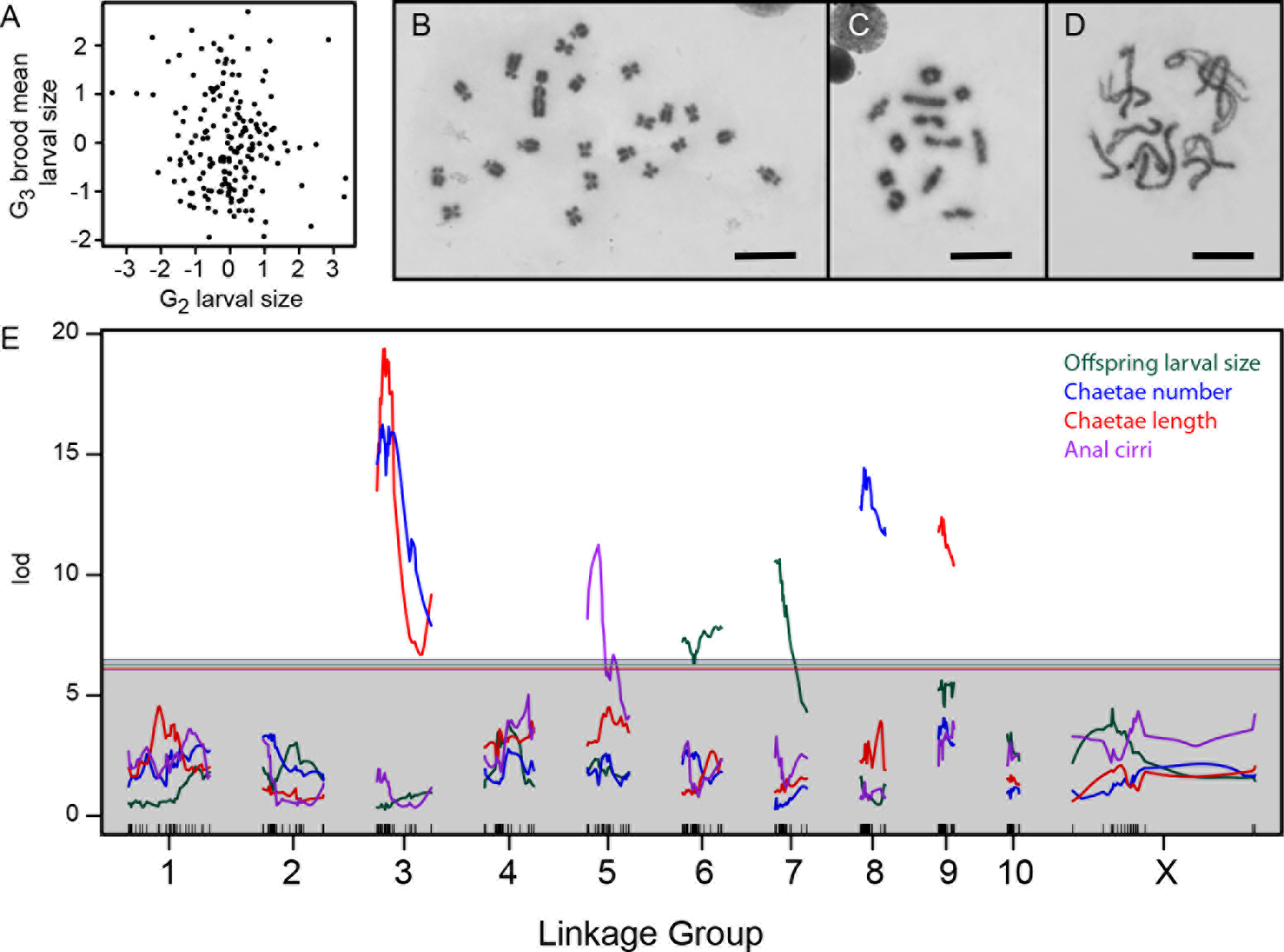
Genetics of developmental mode. **A.** Larval size is uncorrelated between the G_2_ and G_3_ generations (*r*^2^ = −0.1, p = 0.16), consistent with maternal inheritance. **B.** The 22 chromosomes of a male from the Bayonne population include a large Y. Scale bar is 10 μm in panels B-D. **C.** Meiotic chromosomes at diakinesis in the same animal show 12 objects, putatively 10 autosomal bivalents and two sex-chromosome univalents. **D.** Zygotene in a Bayonne female shows 11 bivalents. **E.** Interval mapping of four traits in the G_2_ population identifies seven linkages on six chromosomes. Thresholds for genome-wide significance at p=0.05 are indicated for each trait by horizontal lines, determined by structured permutations that account for G_2_ family structure.

Next we constructed a genetic map, the first for any annelid (File S1). We genotyped F_1_ and G_2_ animals at 702 markers, which fell into 10 autosomal and one sex-linked linkage group (LG)^39,40^. In parallel, we defined the karyotype for *S. benedicti*, which includes 10 pairs of autosomes and one pair of sex chromosomes (Fig 2b). Males are hetorogametic, diagnosed by a large Y chromosome. The karyotypes are grossly indistinguishable between Long Beach and Bayonne, and in each population we observe 11 bivalents at meiotic prophase in females and 12 bodies, presumably 10 autosomal bivalents and 2 sex-chromosome univalents, at diakinesis in males (Fig 2c, d).

G_2_ larvae are intermediate in size and variable in morphology; 15% lack larval chaetae and 25% lack anal cirri. As expected for a maternal trait, G_2_ larval size showed no linkage to G_2_ genotype. Every other phenotype implicated a small number of large-effect loci (Fig 2e). Moreover, the loci are almost entirely independent: the maternal-effect loci shaping larval size are unlinked to the zygotic-effect loci shaping larval morphology. The morphology loci are also largely independent, with a major-effect anal cirri locus on LG5 and loci uniquely affecting number and length of swimming chaetae on LG8 and LG9. The two chaetal traits also share a common locus on LG3. Each QTL has a large effect (Fig 3a) and explains a large proportion of trait variance (Table 1), and the effects are largely additive (Table S1). The two exceptions to additivity are in chaetae length at the LG3 locus (the chaetae-shortening allele is dominant) and larval size, where there is significant epistasis between the two maternal-effect loci on LG6 and LG7. We corroborated the single-trait mapping results by performing a fully multivariate mapping analysis for maternal-effect larval size, chaetae number, and chaetae length. This analysis identified the same loci as the univariate analyses, and showed that the maternal effects and zygotic effects are largely independent (i.e., effects are nearly orthogonal in this three-dimensional space: Fig 3b).

**Figure 3.**
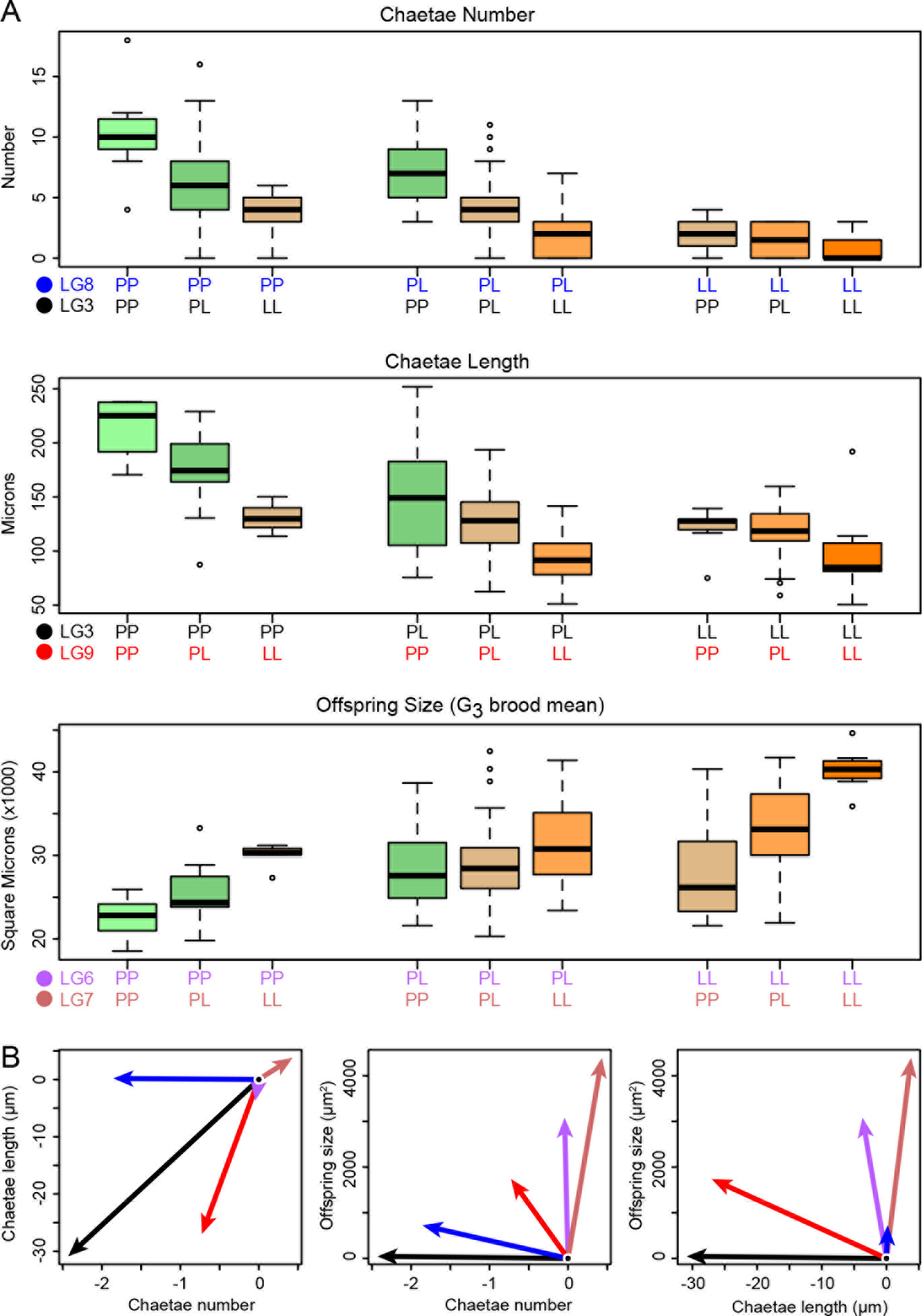
Phenotype distributions for each genotype. **A.** Loci have substantial and largely additive effects. Each boxplot shows the phenotype distribution as a function of the specified G_2_ two-locus genotype, and each box is colored according to the number of planktotroph (P) and lecithotroph (L) alleles it carries. **B.** Multivariate analysis shows that the effects of the five loci in panel A are largely restricted to maternal or zygotic traits. The additive-effect vectors for lecithotrophic alleles in three-trait space are here projected onto two dimensions. Arrow color corresponds to the locus colors in panel A. The major maternal-effect loci are strikingly nearly orthogonal to the major zygotic effects (LG3 and LG8), while the locus on LG9 is mildly pleiotropic, with major effects on chaetae length and minor effects on number and offspring size (all in the aligned direction: shorter chaetae, fewer chaetae, larger offspring).

**Table 1.** G_2_ phenotypic variance explained by significant loci and interactions.

| Trait | Locus | Variance Explained* |
| --- | --- | --- |
| Number of chaetae | LG3 | 21.8% |
|  | LG8 | 18.2% |
| Length of chaetae | LG3 | 25.8% |
|  | LG9 | 13.5% |
| G<sub>3</sub> mean larval size | LG6 | 18.8% |
|  | LG7 | 25.4% |
|  | LG6 x LG7 | 5.2% |
| Presence of anal cirri | LG5 | 18.9% |

\*Percent variance explained is estimated by dropping the specified locus or interaction from the best-fitting genetic model for the phenotype (File S4). In the case of anal cirri, the reported number is the percent deviance explained in a logistic regression.

Our results imply that matings between animals of alternative developmental modes will routinely generate mismatches between larval size and morphology. This genetic architecture creates a strong selective barrier to adaptation: zygotic effect alleles that migrate into new populations will find themselves tested against the local maternal-effect background^20,33,41^. This maternal-by-zygotic epistasis for fitness can inhibit adaptation to local environmental conditions by creating a positive frequency dependence: the locally common maternal background creates the selective regime for zygotic-effect alleles. However, this model will only apply if the zygotic-effect alleles are penetrant across maternal-effect backgrounds. An alternative is that zygotic effects are masked in mismatched maternal backgrounds, allowing them to persist at low frequencies in populations. To test this possibility, we backcrossed 30 G_2_ males (which have an F_1_ maternal background) to planktotrophic females, thereby shifting the maternal background to assess the penetrance of the chaetae loci. We found that the zygotic-effect loci that affect chaetae length and number in an F_1_ maternal background do so in the planktotroph maternal background as well (length: p < 10^−4^; number p < 10^−6^; Fig 4). We estimate a negligible effect of the locus on LG9, but our analysis of that locus is underpowered due to unbalanced genotype frequencies among the 30 G_2_ males.

**Figure 4.**
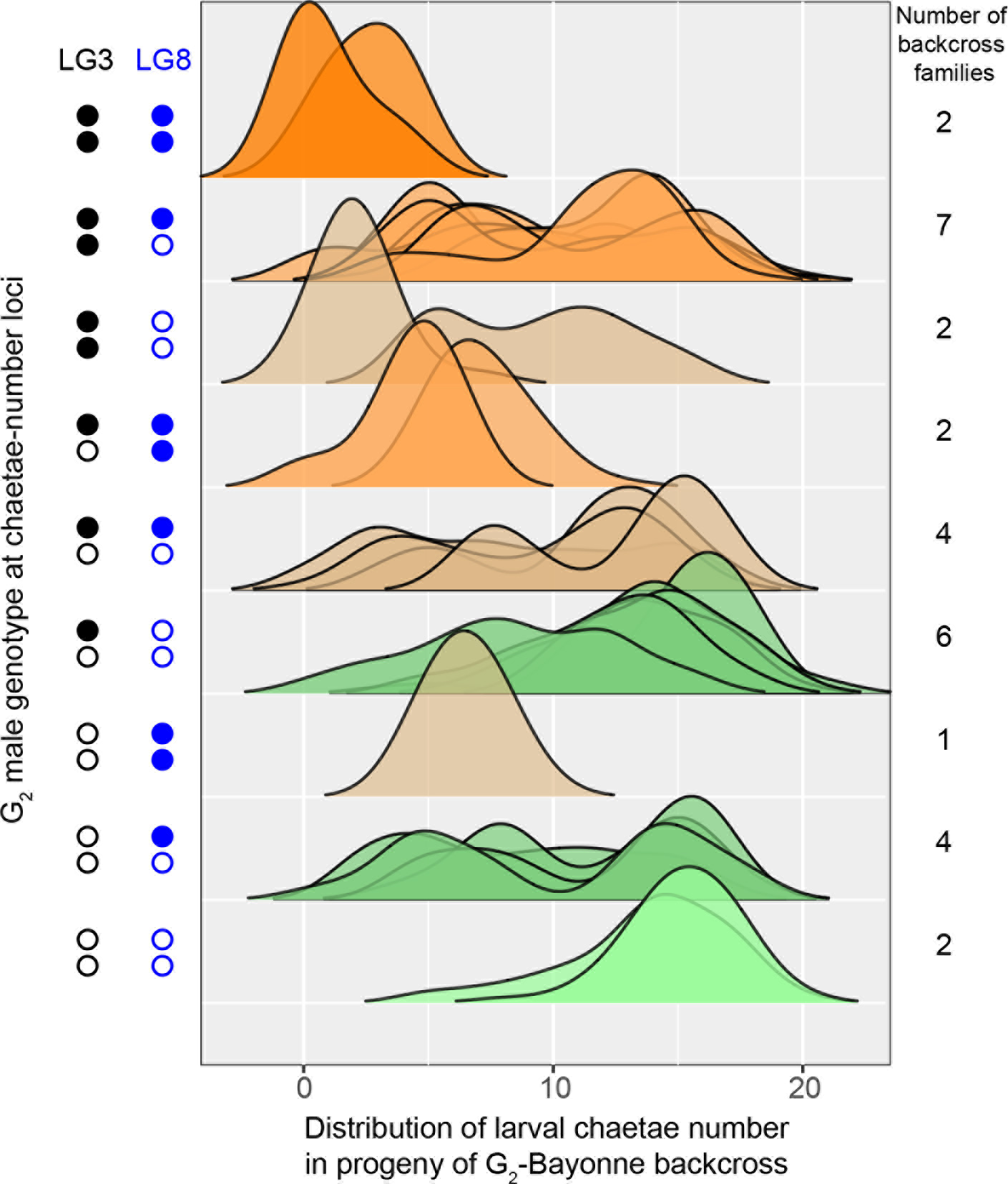
Zygotic alleles are penetrant in a planktotrophic maternal-effect background. Each density shows the distribution of chaetae number in a family derived from a cross between a G_2_ male and a planktotrophic female from the Bayonne population. The genotypes of the G_2_ males are shown at left (filled symbols represent lecithotrophic alleles), and the densities are colored as in Figure 3a by the number of planktotroph and lecithotroph alleles. Note that the progeny of heterozygous G_2_ males have multimodal distributions, demonstrating segregation of large-effect alleles.

The backcross data demonstrate that zygotically-acting alleles are penetrant in planktotrophic maternal backgrounds, and thus these alleles cannot fully “hide” in mismatched populations. However, it remains possible that variation within genotypic classes is such that an occasional larva will have a fully planktotrophic or lecithotrophic phenotype despite carrying mismatched alleles^19^. This raises the possibility that maternal effects can allow for the accumulation of standing genetic variation in a population by masking unfit genotypic combinations from strong selection. Moreover, maternal-effect loci are sex-limited in their expression, and alleles can thus persist in populations at higher frequencies than unconditionally expressed alleles.

We find that co-adapted life-history traits are strikingly modular, where each of the phenotypes has an independent genetic basis with QTL occurring on different chromosomes. This suggests the lecithotrophic larval strategy evolved through the stepwise accumulation of multiple genetic changes throughout the genome. Maternally-determined larval size has an independent genetic basis from other traits that are perfectly correlated in natural populations. While the most direct evolutionary path from plantkotrophy to lecithotrophy involves pleiotropic mutations that move populations diagonally across the size/shape space (Fig 1b), we observe that mutational effects are at right angles to one another. The lack of strong correlated effects among alleles is compatible with the evolution of lecithotrophy via a facultatively planktotrophic intermediate; that is, large larval size may have evolved prior to the loss of larval morphology (Fig 1b).

Although the loci we detect could each represent a cluster of genes, they are located on different chromosomes, indicating that correlated life-history phenotypes are genetically separable. We identified six chromosomes with large-effect alleles for four traits, implying that *S. benedicti* populations can segregate 3^6^ = 729 possible genotypic combinations, most of which are maladaptive. Natural populations consist of only one of the two phenotypic extremes, often occurring in close geographic proximity but rarely directly overlapping^17,28,42^. Nevertheless, Long Beach and Bayonne populations exhibit little genetic differentiation^37,38^, and natural populations that differ in developmental mode show evidence for ongoing gene flow^28^.

Finally, we found that the evolutionary transition to lecithotrophic development requires the interaction of two genotypes, the mother and offspring. The fitness of a zygotic allele depends on the probability it occurs in a favorable maternal genetic background. This genetic architecture provides an explanation for the phenotypic homogeneity of natural populations. The independent genetic basis of maternal and zygotic genetic effects on development shapes the accumulation of variation necessary for selection and local adaptation.

## Acknowledgements

We would like to thank D. Tandon, I. Ukegbu, R. Freih, C. Fayyazi, and L. Jessell for laboratory assistance with maintenance and crossing of the animals; L. Noble, M. Bernstein, J. Yeun for helpful discussions of the manuscript. We thank B. Pernet for collecting the Long Beach worms. This research is funded by the Zegar Family Foundation, NSF grant I0S-1350926 to MVR, NIH grant GM108396-02 to CZ, and an NYU Biology Master’s Research Grant to ADK.

## Supplemental Files

**File S1**: R workspace file containing *S. benedicti* genetic maps and G_2_ phenotype data.

**File S2**: ․csv file containing phenotype and genotype data for Bayonne backcross larvae.

**File S3**. Directory containing data and scripts used to generate the *S. benedicti* genetic maps presented in File S1.

**File S4**. R script file containing the annotated workflow underlying all genetic mapping and phenotypic analyses presented in the manuscript.

## Supplemental methods and results

### 1. Cross Design

We crossed a single outbred Bayonne planktotrophic female to a Long Beach lecithotrophic male to generate F_1_s. Four F_1_ females were crossed to a single F_1_ male to produce 4 half-sib G_2_ families. We intercrossed the G_2_ animals to generate G_3_ families, and we crossed a subset of G_2_ males to Bayonne females to generate backcross families.

### 2. Phenotyping

We imaged G_2_ individuals live so they could be reared to adulthood and crossed to produce G_3_s. A panel of 264 G_2_ larvae was collected within 24 hours of release from their mother and pipetted individually onto a 0.2% agar-seawater pad on a depression slide. We imaged larvae with a Zeiss Axio Imager M2 with 20x DIC objective, and washed them into individual wells in a 24-well plate where they developed to juvenile stage, provisioned with *Dunaliella salina* algae. We measured larval size (area), chaetae number, length of the longest chaeta, and presence/absence of pronounced anal cirri from images using ImageJ (NIH).

Once G_2_ animals reached adulthood, fed with defaunated mud, we paired them randomly for crosses. Because offspring size is proportional to egg size and has a strong additive maternal basis ^19^, we use average G_3_ larval size as a phenotype of the G_2_ mother. G_3_ larvae were fixed in 4% formaldehyde overnight, transferred to PBS buffer, and mounted on slides in a long-term stable mountant. We then imaged and measured larval area as above. Fixed larvae maintained their morphology relative to live larvae when compared from the same clutch. Backcross larvae were fixed and measured in the same manner.

Raw phenotype data are included as part of an Rqtl cross object in File S1, and data for backcross larvae are included in File S2.

### 3. Genotyping

For each Parent, F_1_ and G_2_ individual, we extracted DNA using the Qiagen DNeasy kit. Total worm DNA was sent to SNPsarus, LLC for NextRAD genotyping-by-sequencing. NextRAD uses a primer-based approach to amplify a selection of sequences throughout the genome in order to have a reduced representation library for sequencing. PCR primers amplify genomic loci consistently between samples^53^.

376 libraries were sequenced on Illumina HiSeq 2500 to a depth of ~20x coverage. Reads were aligned using Samtools^54^ to a 805MB (in fragments over 300bp) reference genome assembly. The reference genome was constructed from Illumina Hiseq PE-100 reads of three male *S. benedicti* libraries (insert sizes of 300, 500, 800bp) assembled with Platanus^43^.

SNPsarus aligned reads to the reference genome at 97% identity to reduce spurious alignments, and allowed only two mismatches per read to the reference. SNPs were called in vcftools^44^ and filtered to meet QC requirements by removing loci that had more than two alleles or less than 10% occurrence across all samples. From more than a million scaffolds in the reference assembly, 939 contained SNPs that passed QC, resulting in 1,389 SNPs.

### 4. Karyotyping

We imaged giemsa-stained chromosomes from cells at mitotic metaphase from laboratory-raised inbred (F_5_-F_8_) males and females from the Bayonne and Long Beach populations. We also imaged meiotic diakinesis in males and zygotene in females. We used a karyotyping protocol modeled closely on Rowell et al^45^. Briefly, worm tissue was incubated in 0.05% colchicine in artificial seawater for 90 minutes, then subjected to a hypotonic series reaching 1:4 ASW:H_2_0 over one hour. The tissue was then fixed with fresh Carnoy’s Fixative, with three fixative changes over one hour. Next, tissue was macerated on a glass slide in 60% acetic acid, using glass needles. Finally, after the slides had dried, we stained them with 5% Giemsa stain in phosphate-buffered saline. We imaged the slides on a Zeiss Axio Imager M2 with 100x objective.

Preparations from both sexes from both populations revealed a consistent karyotype of 2n=22 and no gross differences between populations. Males are heteromorphic, carrying a distinctive Y chromosome that is substantially larger than every other chromosome. Among annelids with XY chromosomal sex determination, the Y is often larger than the X, as in species of *Hediste^46^*. In *Polydora curiosa*, the only spionid previously known to have heteromorphic sex chromosomes (2n=34XY), the Y is larger than all other chromosomes^47^, as we observe in *S. benedicti*. Many other species of *Polydora* lack heteromorphic sex chromosomes^47^, leaving the potential homology of the large Y chromosomes in *P. curiosa* and *S. benedicti* uncertain.

Meiotic material from females of each population shows 11 bivalents at zygotene. In male meotic material from each population, we observe 12 structures at diakinesis, including a mixture of ring and rod bivalents. We hypothesize that two of the 12 structures are X- and Y-chromosome univalents. This pattern of unpaired X and Y chromosomes at metaphase I is known among plant and insect species^48^. Korablev et al.^47^ observed that the X and Y pair end-to-end at metaphase I in *Polydora curiosa*, which therefore differs from *S. benedicti*.

### 5. Linkage Map construction

Data and annotated scripts used to generate *S. benedicti* linkage maps are included in File S3.

We filtered data to include only the 313 unique samples for which the mean read depth at the 1,389 SNP markers was greater than 15 and for which more than 1,000 SNP genotypes were called. Unfortunately, due to sample quality issues, we were unable to generate reliable genotype calls from the two P_0_ animals or from the F_1_ male and four F_1_ females that parented the four F_2_ families. We retained 47 F_1_ and 266 G_2_ individuals.

We tested each SNP for sex linkage by applying Fisher’s exact test to genotype counts within each family and then combining the p-values using Fisher’s meta-analysis. For non-sex-linked contigs that contain multiple SNPs, we manually recoded each contig, when possible, to be a single intercross marker, with inferred P_0_ genotypes AA × BB. These inferences are straightforward given multiple SNPs per contig and our data from a panel of F_1_s and four half-sib G_2_ families. In some cases, individual G_2_ families segregated haplotypes that were not all distinguishable. For example, if P_0_ haplotypes were AB x AC, segregation in G_2_ families descended from AC × BC F_1_ parents is fully resolvable, but segregation from AA × BC F_1_ parents is not, because the P_0_ origin of A haplotypes in the G_2_ is ambiguous. In such cases, we assigned partial genotypes in the relevant families (i.e., we treated them as dominant rather than codominant markers). However, we only recoded multi-SNP contigs for which segregation in at least one G_2_ family was fully resolvable. Note that the assignment of genotypes to the maternal vs. paternal P_0_ is arbitrary because we lack genotypes for those individuals. To call a marker genotype for an individual worm, we required that all SNPs in the contig marker had to have genotype calls that conform to an intact P_0_ haplotype; this adds a layer of stringency in excluding genotypes that contain any erroneous SNP calls. Most multi-SNP contigs contain two or three SNPs, but one contained 16 SNPs spread across multiple sequencing reads. Because of missing data and genotype error, the requirement for intact P_0_ haplotypes across 16 SNPs is too stringent, and this contig was recoded into 9 separate markers (five multi-SNP and four single SNP, based on the distribution of the SNPs across sequencing reads).

In recoding multi-SNP contigs as single markers, we identified five strains whose data showed very high rates of incompatibility with possible segregation patterns and 16 strains that showed probable pedigree errors (i.e., their genotypes were inconsistent with maternity by the F_1_ mother shown in their records but compatible with one of the other F_1_ mothers). These 21 strains were excluded from subsequent analysis, leaving a dataset of 292 worms distributed across five families:

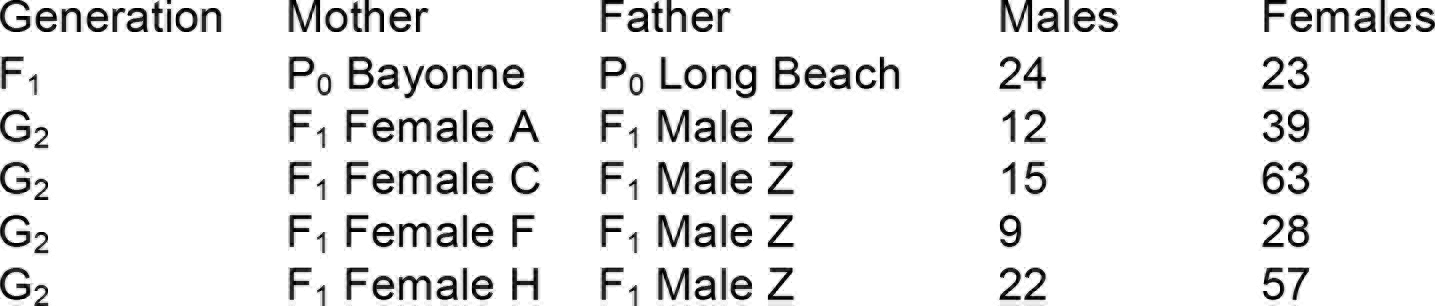
Number of individuals from each family used in mapping crosses.

We inferred segregation classes for each autosomal marker and imputed genotypes for the five parents of the G_2_ families using a composite likelihood approach. For every autosomal SNP, there are four possible P_0_ genotype combinations (AAxBB, ABxAB, AAxAB, ABxBB), plus two for sites erroneously called as SNPs (AAxAA, BBxBB). For each SNP, we estimated likelihoods for each configuration for each family. We maximized the likelihood including a one-step error probability (i.e., AA <-> AB <-> BB) allowed to range up to 10%. We next identified the segregation pattern (including the genotypes of the G_2_ parents of the G_2_ families) that maximizes the likelihood over the five families. This is a composite likelihood insofar as the genotype error probabilities are estimated for each family and the global pedigree likelihood derived from the appropriate combination of family likelihood estimates.

We then used three steps to build the *S. benedicti* genetic map:

1. Construction of an autosomal map from markers segregating in an intercross pattern, using Rqtl ^39^.
2. Construction of an autosomal map using the refined intercross marker dataset from step 1 and hundreds of additional non-intercross autosomal markers, using lep-MAP _49,50_.
3. Addition of a sex-linked linkage-group.

#### Step 1: Intercross marker map

The first strategy uses data from 549 non-sex-linked SNPs that individually or in contig-groups segregate in an intercross pattern (i.e., F_1_s are heterozygous and G_2_s segregate 1:2:1, implying that the parental genotypes are homozygous for alternate alleles). These SNPs include 448 from multi-SNP contigs that are recoded into 193 intercross markers, plus 101 individual SNPs identified as intercross by likelihood. We excluded 17 markers that were missing data for more than 10% of individuals and 3 other markers that exhibited strong departures from Mendelian proportions (p < 10^−10^). From the resulting dataset of 274 markers and 245 G_2_ individuals, we constructed an autosomal genetic map in Rqtl (v.1.40-8) following the protocol of Broman^51^. 271 markers readily assembled into 10 autosomal linkage groups. We ordered the markers using 20 cycles of the *orderMarkers* function with ripple window = 7, retaining the ordering with the highest likelihood for each linkage group. Four worms exhibited a large excess of crossovers, indicative of high genotyping error rates. We removed these animals (all are Family C females), estimated the genotyping error rate as 0.013 by maximum likelihood, and reordered the markers with another 20 cycles of *orderMarkers* with error.prob = 0. 013.

Some apparent crossovers are associated with single genotypes that differ from both immediate flanking markers, despite very small genetic distances. These represent probable genotyping errors. Using an error lod of 6 as a threshold for probable errors, we converted 420 (of almost 63,000) genotype calls to missing data. We then reestimated the maximum likelihood error rate as 0.004 for the remaining data and reordered the markers with 50 cycles of *orderMarkers* and error.prob = 0.004.

#### Step 2. Inclusion of non-intercross markers

Returning to the original 1,389-SNP dataset, we replaced the intercross marker genotypes with those from step 1 (i.e., replacing multi-SNP contig SNPs with their inferred intercross gentoypes, converting 420 genotypes to missing data, and removing the four individuals inferred to have high genotype error rates). For the remaining non-intercross SNPs, we imputed the genotypes of the five F_1_ parents by likelihood, as described, and we used lepmap2^49^ (v.0.2, downloaded March 5, 2017) to assign markers to linkage groups. The input dataset includes 288 individuals (47 F_1_s and 241 G_2_s in four families) and 1,111 markers. Processing steps included *ParentCallGrandparents* with XLimit=8 and ZLimit=8, *Filtering* with epsilon=0.02 and dataTolerance=0.0001, *SeparateChromosomes* with lodLimit=15, and *JoinSingles* with lodLimit=10 and lodDifference=5. This analysis assigned 676 markers to 10 autosomes and 97 markers to two sex-linked chromosomes (Fig 2). An additional 25 markers were assigned to 10 small linkage groups (one sex-linked) with two to four markers each, and 313 markers were not assigned to linkage groups. The ten major autosomal linkage groups mapped uniquely to the ten identified using the intercross markers above at step 1.

To order the autosomal markers, we used *OrderMarkers2* in lepmap3^50^ (v.0.1, downloaded March 5, 2017). For each linkage group, we ran 100 iterations with randomPhase=0 and 100 iterations with randomPhase=1, and we then repeatedly ran *OrderMarkers2* with the evaluateOrder option on the highest likelihood results for each approach to search for higher likelihood reconstructions. We then selected the result with the highest likelihood overall and retained the reconstructed genotypes (outputPhasedData=1) for that run.

We next converted these reconstructed genotypes to conventional intercross genotypes by mapping the intercross-marker genotypes onto them and retaining the phase with the highest likelihood. We renamed the linkage groups to sort them by genetic map length and the result is **Sb676**. The intercross map from step 1, with linkage groups named accordingly, is **Sb271**.

#### Step 3. Sex-linked markers

For each of the 1,389 SNPs in the original dataset, we estimated the likelihood of each possible autosomal or sex-linked segregation pattern. We identified 32 SNPs with clean X-linked segregation (AAxBY or BBxAY) and none with Z-linkage. The X-linked SNPs mapped to one major and one minor sex-linked linkage group in the lepmap analysis at step 2 above. The other major lepmap-defined sex-linked linkage group contained only SNPs that segregate as fixed differences between X and Y chromosomes (i.e., the maximum likelihood segregation patterns are ^X^A^X^A x ^X^A^Y^B and ^X^B^X^B x ^X^B^Y^A). These are uninformative for the genetic map.

We next examined the multi-SNP contigs that contain any SNP whose sex-linkage p-value is less than 0.01, and where possible we recoded these SNPs as single multi-SNP markers, converting impossible genotypes to missing data, as done for autosomal markers in step 1. This resulted in a total of 26 markers (10 multi-SNP and 16 single-SNP). These markers were then ordered in Rqtl as in step 1. A group of three markers, which form a separate, minor sex-linked linkage group in our lepmap analysis at step 2, here form a distantly linked piece of the X chromosome (min.lod = 6, max.rf=0.33). The final map, incorporating 702 markers, is **Sb702**.

The maps produced at each of the three steps are provided as Rqtl cross objects in File S1.

### 6. QTL Mapping

We performed interval mapping in Rqtl to identify QTL. To accommodate the family structure in the data, we performed genome scans separately in each G_2_ family and summed the lod scores. We determined the genome-wide significance by following the same strategy after permuting the data within G_2_ families. All traits were mapped assuming a normal model, except for the presence or absence of anal cirri, for which we used a binary (logistic) model. R scripts that reproduce all linkage mapping and phenotypic analyses in the paper are included in File S4.

To model the genetics of each trait using the Rqtl *fitqtl* function, we tested for pairwise interactions between detected QTL across linkage groups and for effects of G_2_ family. In each case we compared a model of additive and dominant within-locus effects to models that incorporated the additional terms, and we used likelihood ratio tests to determine the statistical significance of improvements in model fit. We estimate the effect size for QTLs using the best-fitting model, as described in File S4. For chaetae number and presence of anal cirri, the prefered model includes QTL effects only. For chaetae length, the model includes an effect of G_2_ family. For G_3_ offspring area, the model includes an interaction between two QTL. Effect estimates are reported below in Table S1.

**Table S1.**
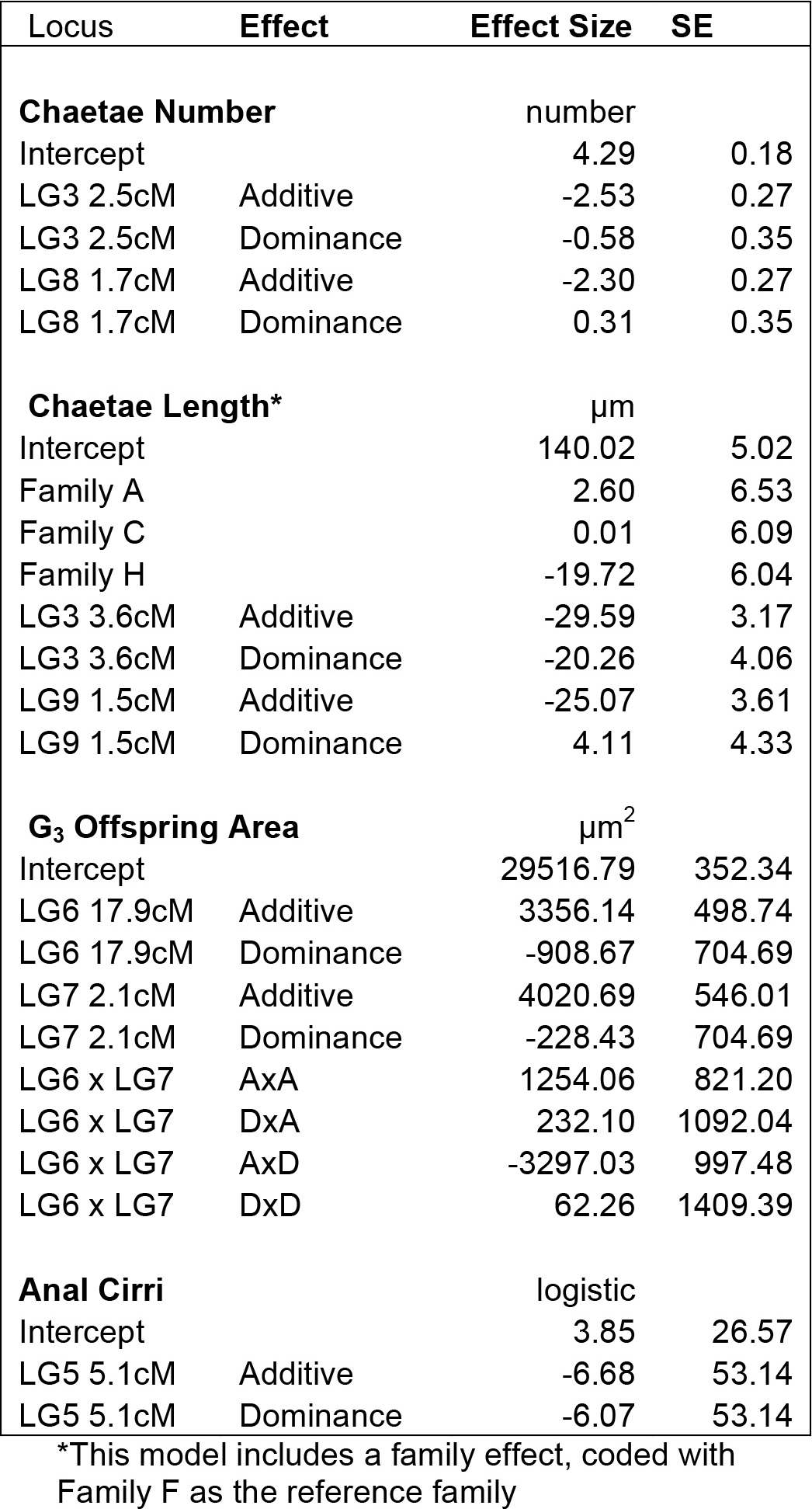
Effect sizes for significant QTL for each trait.

We find that chaetae number and length are correlated, as in ref ^19^, (*r*^2^ = 0.22, p<0.001). The two traits share a QTL on LG3 that may act pleiotropically. However, when variation due to the LG3 QTL is removed, there is still a strong correlation between the trait residuals (*r*^2^ = 0.1, p<0.001).

### 7. Multivariate QTL scan

We analyzed the three continuous traits - mean offspring size, chaetae number, and chaetae length - in a fully multivariate QTL scan^52^. The approach searches for regions of the genome with significant effects in the three-dimensional space defined by variation in these three traits. We restricted the dataset to the 149 G_2_ females for which all three traits were scored (i.e., excluding G_2_ worms with zero chaetae).

We fit the following model at each marker: **Y = XB + e**, where **Y** is the 149x3 matrix of phenotypes, **X** is a matrix of fixed effects, **B** is the vector of fixed-effect coefficients to be estimated, and **e** is normally-distributed residual error. The fixed effects are the intercept, G_2_ family identity, and two vectors coding the additive and dominance variables for a single genetic marker. At each marker, we fit this model and compared it to a model with intercept and G_2_ family as the only fixed effects, calculating a p-value from the Pillai-Bartlett statistic. We estimated thresholds for genome-wide significance by repeating the analysis on 1000 datasets in which phenotype vectors were permuted among individuals within G_2_ families. R code that reproduces the analysis is included in File S4.

Five QTL exceeded the genome-wide threshold for significance (p ≤ 0.001 in each case), coinciding with the five QTL detected for these traits by univariate analysis. In a model incorporating additive and dominance effects for each QTL, five additive and two dominance effects (for the QTL on LG3 and LG6) were retained as significant (p < 0.01; Table S2). We then estimated QTL effect sizes under the reduced model that includes only the significant effects.

**Table S2.**
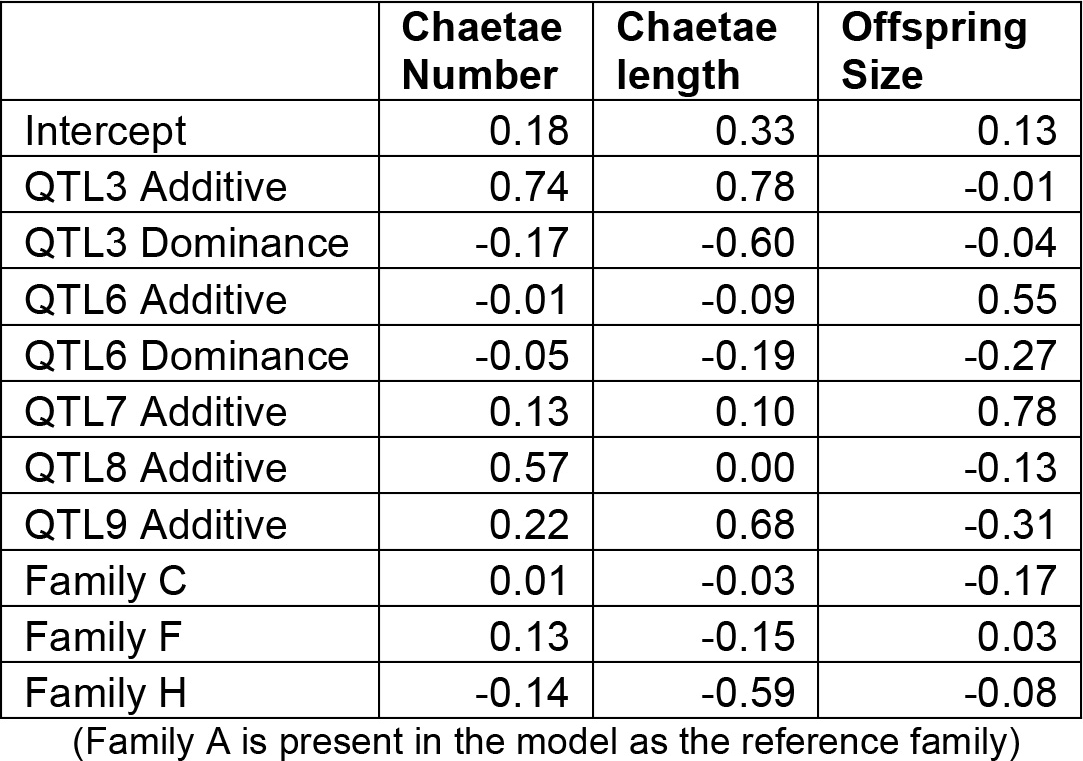
Estimated effects from reduced multivariate model QTL scan (in units of G_2_ phenotypic standard deviations)

### 8. Analysis of backcross larvae

We tested for an effect of G_2_ male genotype on the distribution of chaetal phenotypes in their backcross progeny. We used a linear mixed-effect model to account for the relatedness of larvae within a brood and then compared models with and without the loci implicated in each trait in the G_2_ generation (i.e., the loci in Table S1). For both traits, chaetal length and number, we detected a significant effect of genotype, with effects in the expected directions. The effect size estimates indicate that the locus on LG9 does not have a significant effect on chaetal length. We note that the 30 G_2_ males used in the backcross have unbalanced genotype representation at this locus, with 8 LL homozygotes, 22 LP heterozygotes, and 0 PP homozygotes. We are therefore poorly powered to detect an effect of this locus. R code reproducing our analyses are included in File S4.

## References

1 Collin, R. Phylogenetic effects, the loss of complex characters, and the evolution of development in calyptraeid gastropods. Evolution 58, 1488–1502, (2004).

2 Hart, M. W., Byrne, M. & Smith, M. J. Molecular phylogenetic analysis of life-history evolution in Asterinid starfish. Evolution 51, 1848–1861, (1997).

3 Smith, M. S., Zigler, K. S. & Raff, R. A. Evolution of direct-developing larvae: selection vs loss. Bioessays 29, 566–571, (2007).

4 Strathmann, R. R. Feeding and nonfeeding larval development and life-history evolution in marine-invertebrates. Annual Review of Ecology and Systematics 16, 339–361, (1985).

5 Thorson, G. Reproductive and larval ecology of marine bottom invertebrates. Biological Reviews of the Cambridge Philosophical Society 25, 1–45, (1950).

6 Wray, G. A. Parallel evolution of nonfeeding larvae in echinoids. Systematic Biology 45, 308–322, (1996).

7 Raff, R. A. The shape of life: genes, development, and the evolution of animal form. (University of Chicago Press, 1996).

8 Christiansen, F. B. & Fenchel, T. M. Evolution of marine invertebrate reproductive patterns. Theoretical Population Biology 16, 267–282, (1979).

9 Vance, R. R. Reproductive strategies in marine benthic invertebrates. American Naturalist 107, 339–352, (1973).

10 Vance, R. R. More on reproductive strategies in marine benthic invertebrates. American Naturalist 107, 353–361, (1973).

11 Havenhand, J. N. in Ecology of marine invertebrate larvae (ed L. R. McEdward) 79–122 (CRC Press, 1995).

12 Arndt, A. & Smith, M. J. Genetic diversity and population structure in two species of sea cucumber: differing patterns according to mode of development. Molecular Ecology 7, 1053–1064, (1998).

13 Duda, T. F. & Palumbi, S. R. Developmental shifts and species selection in gastropods. Proceedings of the National Academy of Sciences of the United States of America 96, 962–10277, (1999).

14 Jeffery, C. H. & Emlet, R. B. Macroevolutionary consequences of developmental mode in temnopleurid echinoids from the tertiary of southern Australia. Evolution 57, 1031–1048, (2003).

15 McEdward, L. R. & Miner, B. G. Larval and life-cycle patterns in echinoderms. Canadian Journal of Zoology-Revue Canadienne De Zoologie 79, 1125–1170, (2001).

16 Wray, G. A. in Ecology of marine invertebrate larvae (ed L. R. McEdward) 413–447 (CRC Press, 1995).

17 Levin, L. A. Multiple patterns of development in Streblospio benedicti Webster (Spionidae) from 3 coasts of North America. Biological Bulletin 166, 494–508, (1984).

18 Levin, L. A., Zhu, J. & Creed, E. The genetic basis of life-history characters in a polychaete exhibiting planktotrophy and lecithotrophy. Evolution 45, 380–397, (1991).

19 Zakas, C. & Rockman, M. V. Dimorphic development in Streblospio benedicti: genetic analysis of morphological differences between larval types. International Journal of Developmental Biology 58, 593–599, (2014).

20 Wolf, J. B. & Brodie, E. D. The coadaptation of parental and offspring characters. Evolution 52, 299–308, (1998).

21 Blake, J. A. Reproduction and larval development of Polydora from northern New England (Polychaeta Spionidae). Ophelia 7, 1–63, (1969).

22 Gibson, G., MacDonald, K. & Dufton, M. Morphogenesis and phenotypic divergence in two developmental morphs of Streblospio benedicti (Annelida, Spionidae). Invertebrate Biology 129, 328–343, (2010).

23 McCain, E. R. Poecilogony as a tool for understanding speciation: Early development of Streblospio benedicti and Streblospio gynobranchiata (Polychaeta:Spionidae) (vol 51, pg 91, 2008). Invertebrate Reproduction & Development 52, 121–121, (2008).

24 Pernet, B. & McHugh, D. Evolutionary changes in the timing of gut morphogenesis in larvae of the marine annelid Streblospio benedicti. Evol Dev 12, 618–627, (2010).

25 Levin, L. A. & Bridges, T. S. Control and consequences of alternative developmental modes in a poecilogonous polychaete. American Zoologist 34, 323–332, (1994).

26 Levin, L. A. & Creed, E. L. Effect of temperature and food availability on reproductive responses of Streblospio benedicti (Polychaeta, Spionidae) with planktotrophic or lecithotrophic development. Marine Biology 92, 103–113, (1986).

27 Schulze, S. R., Rice, S. A., Simon, J. L. & Karl, S. A. Evolution of poecilogony and the biogeography of North American populations of the polychaete Streblospio. Evolution 54, 1247–1259, (2000).

28 Zakas, C. & Wares, J. P. Consequences of a poecilogonous life history for genetic structure in coastal populations of the polychaete Streblospio benedicti. Molecular Ecology 21, 5447–5460, (2012).

29 Whitlock, M. C., Phillips, P. C., Moore, F. B. G. & Tonsor, S. J. Multiple fitness peaks and epistasis. Annual Review of Ecology and Systematics 26, 601–629, (1995).

30 Hall, M. C., Basten, C. J. & Willis, J. H. Pleiotropic quantitative trait loci contribute to population divergence in traits associated with life-history variation in Mimulus guttatus. Genetics 172, 1829–1844, (2006).

31 Jones, F. C. et al. The genomic basis of adaptive evolution in threespine sticklebacks. Nature 484, 55–61, (2012).

32 Joron, M. et al. Chromosomal rearrangements maintain a polymorphic supergene controlling butterfly mimicry. Nature 477, 203–U102, (2011).

33 Kirkpatrick, M. & Lande, R. The evolution of maternal characters. Evolution 43, 485–503, (1989).

34 Wolf, J. B. & Wade, M. J. Evolutionary genetics of maternal effects. Evolution 70, 827–839, (2016).

35 Allen, J. D. & Pernet, B. Intermediate modes of larval development: bridging the gap between planktotrophy and lecithotrophy. Evolution & Development 9, 643–653, (2007).

36 Pernet, B. & McArthur, L. Feeding by larvae of two different developmental modes in Streblospio benedicti (Polychaeta : Spionidae). Marine Biology 149, 803–811, (2006).

37 Rockman, M. V. Patterns of nuclear genetic variation in the poecilogonous polychaete Streblospio benedicti. Integrative and Comparative Biology 52, 173–180, (2012).

38 Zakas, C. & Rockman, M. V. Gene-based polymorphisms reveal limited genomic divergence in a species with a heritable life-history dimorphism. Evolution & Development 17, 240–247, (2015).

39 Broman, K. W., Wu, H., Sen, S. & Churchill, G. A. R/qtl: QTL mapping in experimental crosses. Bioinformatics 19, 889–890, (2003).

40 R Core Development Team. R: A language and environment for statistical computing (R Foundation for Statistical Computing, Vienna, Austria, 2015).

41 Wolf, J. B. Gene interactions from maternal effects. Evolution 54, 1882–1898, (2000).

42 Levin, L. A. & Huggett, D. V. Implications of alternative reproductive modes for seasonality and demography in an estuarine polychaete. Ecology 71, 2191–2208, (1990).

43 Kajitani, R. et al. Efficient de novo assembly of highly heterozygous genomes from whole-genome shotgun short reads. Genome Research 24, 1384–1395, (2014).

44 Danecek, P. et al. The variant call format and VCFtools. Bioinformatics 27, 2156–2158, (2011).

45 Rowell, D. M., Lim, S. L. & Grutzner, F. Chromosome analysis in invertebrates and vertebrates. Molecular Methods for Evolutionary Genetics 772, 13–35, (2011).

46 Tosuji, H., Miyamoto, J., Hayata, Y. & Sato, M. Karyotyping of female and male Hediste japonica (Polychaeta, Annelida) in comparison with those of two closely related species, H. diadroma and H. atoka. Zoological Science 21, 147–152, (2004).

47 Korablev, V. P., Radashevsky, V. I. & Manchenko, G. P. The XX-XY (male-heterogametic) sex chromosome system in Polydora curiosa (Polychaeta : Spionidae). Ophelia 51, 193–201, (1999).

48 Brady, M. & Paliulis, L. V. Chromosome interaction over a distance in meiosis. Royal Society Open Science 2, (2015).

49 Rastas, P., Calboli, F. C., Guo, B., Shikano, T. & Merila, J. Construction of ultradense linkage maps with Lep-MAP2: Stickleback F2 recombinant crosses as an example. Genome Biol Evol 8, 78–93, (2015).

50 Rastas, P. Lep-MAP3: robust linkage mapping even for low-coverage whole genome sequencing data. Bioinformatics, btx494, (2017).

51 Broman, K. W. Genetic map construction with R/qtl. Technical Report, Department of Biostatistics and Medical Informatics, University of Wisconsin-Madison 214, (2010).

52 Knott, S. A. & Haley, C. S. Multitrait least squares for quantitative trait loci detection. Genetics 156, 899–911, (2000).

53 Russello, M. A., Waterhouse, M. D., Etter, P. D. & Johnson, E. A. From promise to practice: pairing non-invasive sampling with genomics in conservation. PeerJ, 3, e1106, (2015).

54 Li H., Handsaker B., Wysoker A., Fennell T., Ruan J., Homer N., Marth G., Abecasis G., Durbin R. and 1000 Genome Project Data Processing Subgroup. The Sequence alignment/map (SAM) format and SAMtools. Bioinformatics, 25, 2078–9, (2009).

